# Intracellular Group A *Streptococcus* induces Golgi fragmentation to impair host defenses through Streptolysin O and NAD-glycohydrolase

**DOI:** 10.1101/2020.07.16.207894

**Authors:** Takashi Nozawa, Junpei Iibushi, Hirotaka Toh, Atsuko Minowa-Nozawa, Kazunori Murase, Chihiro Aikawa, Ichiro Nakagawa

## Abstract

Group A *Streptococcus* (GAS; *Streptococcus pyogenes*) is a major human pathogen that causes streptococcal pharyngitis, skin and soft-tissue infections, and life-threatening conditions such as streptococcal toxic-shock syndrome. During infection, GAS not only invades diverse host cells, but also injects effector proteins such as NAD-glycohydrolase (Nga) into the host cells through a streptolysin O (SLO)-dependent mechanism without invading the cells; Nga and SLO are two major virulence factors that are associated with increased bacterial virulence. Here, we have shown that the invading GAS induces fragmentation of the Golgi complex and inhibits anterograde transport in the infected host cells through the secreted toxins SLO and Nga. GAS infection-induced Golgi fragmentation required both bacterial invasion and SLO-mediated Nga translocation into the host cytosol. The cellular Golgi network is critical for the sorting of surface molecules and thus is essential for epithelial barrier integrity and the immune response of macrophages to pathogens. In epithelial cells, inhibition of anterograde trafficking by invading GAS and Nga resulted in the redistribution of E-cadherin to the cytosol and an increase in bacterial translocation across the epithelial barrier. Moreover, in macrophages, interleukin-8 secretion in response to GAS infection was found to be suppressed by intracellular GAS and Nga. Our findings reveal a previously undescribed bacterial invasion-dependent function of Nga as well as a previously unrecognized GAS-host interaction that is associated GAS pathogenesis.

**Importance:** Two prominent virulence factors of GAS, SLO and Nga, have been established to be linked to enhanced pathogenicity of prevalent GAS strains. Recent advances show that SLO and Nga are important for intracellular survival of GAS in epithelial cells and macrophages. Here, we found that invading GAS disrupt the Golgi complex in host cells by SLO and Nga. We showed that GAS-induced Golgi fragmentation requires bacterial invasion into host cells, SLO pore-formation activity, and Nga NADase activity. GAS-induced Golgi fragmentation resulted in the impairment of epithelial barrier and chemokine secretion in macrophages. This immune inhibition property of SLO and Nga by intracellular GAS indicates that the invasion of GAS is associated with virulence exerted by SLO and Nga.

## Introduction

Group A *Streptococcus* (GAS; *Streptococcus pyogenes*) is a human-specific pathogen responsible for diverse diseases, ranging from pharyngitis and impetigo to life-threatening conditions such as necrotizing fasciitis and streptococcal toxic-shock syndrome (STSS) in which mortality rates are 30%–70% even with immediate antibiotic therapy and intensive care [1]. Therefore, GAS species are commonly referred to as “killer bacteria” or “flesh-eating bacteria,” and the ability of GAS to spread rapidly at the infection site and disseminate systemically indicates that the pathogen possesses robust mechanisms to resist the human innate immune response.

The initial sites of GAS infection are the pharyngeal epithelia and the keratinocytes, and the pathogen invades deeper tissues through the paracellular pathway by degrading the junctional proteins. Although GAS is commonly regarded as an extracellular pathogen, GAS can invade epithelial cells, endothelial cells, and macrophages, and this cellular invasion has been reported to be associated with GAS pathogenesis [2-5]. However, several GAS strains excluding certain isolates are degraded through the endosomal pathway or autophagy and cannot survive for long periods inside the epithelial cells [6-11], and the importance of GAS invasion into host cells remains incompletely elucidated.

Recent transcriptome evidence has revealed that highly virulent GAS strains exhibit enhanced expression of two toxins, streptolysin O (SLO) and NAD-glycohydrolase (Nga) [12-14], which emphasizes a role for these toxins in GAS pathogenesis. SLO is a member of the family of cholesterol-dependent cytolysins that bind to the cholesterol-containing membranes, oligomerize, and insert into the lipid bilayer to form pores [15-17]. SLO not only induces the necrosis of neutrophils through pore formation [17,18], but also translocates the effector protein Nga into the host cytosol in a pore-formation independent manner and thereby promotes intracellular survival in macrophages and epithelial cells [19-21]. Nga hydrolyzes NAD into nicotinamide and ADP-ribose and thus depletes intracellular NAD pools and causes ATP depletion in cells. Accordingly, Nga has been reported to inhibit the acidification of phagosomes or autolysosomes potentially through ATP depletion in macrophages and keratinocytes [9,22]. Moreover, Nga inhibits the canonical autophagy pathway to promote bacterial survival in epithelial cells [23], Nga extracellularly inhibits interleukin (IL)-1β production [21], and nicotinamide potently inhibits the secretion of proinflammatory cytokines from monocytes [24]. These lines of evidence have established that SLO and Nga enable GAS to persist within host cells and modulate immune responses, and these effects are considered to be exerted by Nga activity itself.

Pathogenic bacteria evade host defenses by subverting the host signaling pathways through several distinct and sophisticated mechanisms [25,26]. For example, *Legionella pneumophila, Chlamydia trachomatis*, and *Burkholderia thailandensis* secrete the SET-domain-containing proteins that methylate histones to alter the chromatin landscape of the host cell [27-29] and thus promote the intracellular proliferation of bacteria. *Salmonella* Typhimurium, *Legionella* spp., and *Brucella* spp. modulate host membrane dynamics to allow the bacteria to form replication-permissive vacuoles [25]. Enteropathogenic *Escherichia coli* and *Shigella flexneri* target the Golgi network, the ER, and the eukaryotic secretory pathway to suppress the host defenses [30,31]. Here, to uncover the previously unrecognized GAS-host interactions, we examined the organelle morphology in host cells during GAS infection, which revealed that GAS infection triggers the fragmentation of the Golgi complex. We determined that SLO and Nga were responsible for this effect, and further that both SLO-mediated Nga translocation and bacterial invasion into host cells were required for disrupting the Golgi network. Inhibition of the Golgi network resulted in the loss of not only epithelial integrity, but also IL-8 secretion by macrophages in response to GAS infection.

## Results

### Golgi apparatus is fragmented during GAS infection

Because intracellular signaling and vesicular trafficking are closely associated with organelles, disruptions of host functions frequently result in alterations of the organelle morphology, Therefore, we examined the mechanism by which GAS infection affects the intracellular vesicular or signaling networks: We infected the HeLa cells with GAS JRS4 strain, an M6 strain that efficiently invades host cells, immunostained the cells for a series of organelle marker proteins, and examined and compared the morphology of the mitochondria, ER, *cis*-Golgi, and *trans*-Golgi before and after infection. Notably, the infection produced overt morphological changes in the mitochondria and in the *cis*/*trans*-Golgi (Fig. S1). During GAS infection, the normal tubular network of the mitochondria was fragmented into short rods or spheres, and the typical ribbon-like structure of the Golgi complex was also fragmented into punctate structures and dispersed throughout the cytoplasm (Fig. S1). We have previously reported that GAS invasion triggers apoptotic signaling, which causes mitochondrial fragmentation [32]. Thus, in the present study we examined the Golgi fragmentation during GAS infection in more detail. To test whether this infection-induced Golgi fragmentation was observed in different cell types, we used GAS JRS4 for infecting the lung epithelial cells (A549 cells), the human keratinocytes (HaCat cells), the primary dermal keratinocytes (normal human epidermal keratinocytes; NHEKs), the human umbilical vein endothelial cells (HUVECs), and a human monocyte leukemia cell line (THP-1). JRS4 infection fragmented the Golgi structures, which then appeared dispersed throughout the cytoplasm in all types of the cells tested (Fig. 1A). We also infected these cells with two other GAS strains: NIH45, a serotype M28 and an invasive strain isolated from an STSS patient; and 4944, an epidemic serotype M89 clade-3 strain. Both the GAS strains also clearly caused Golgi fragmentation in all the cells examined (Fig. 1A).

**Fig. 1.**
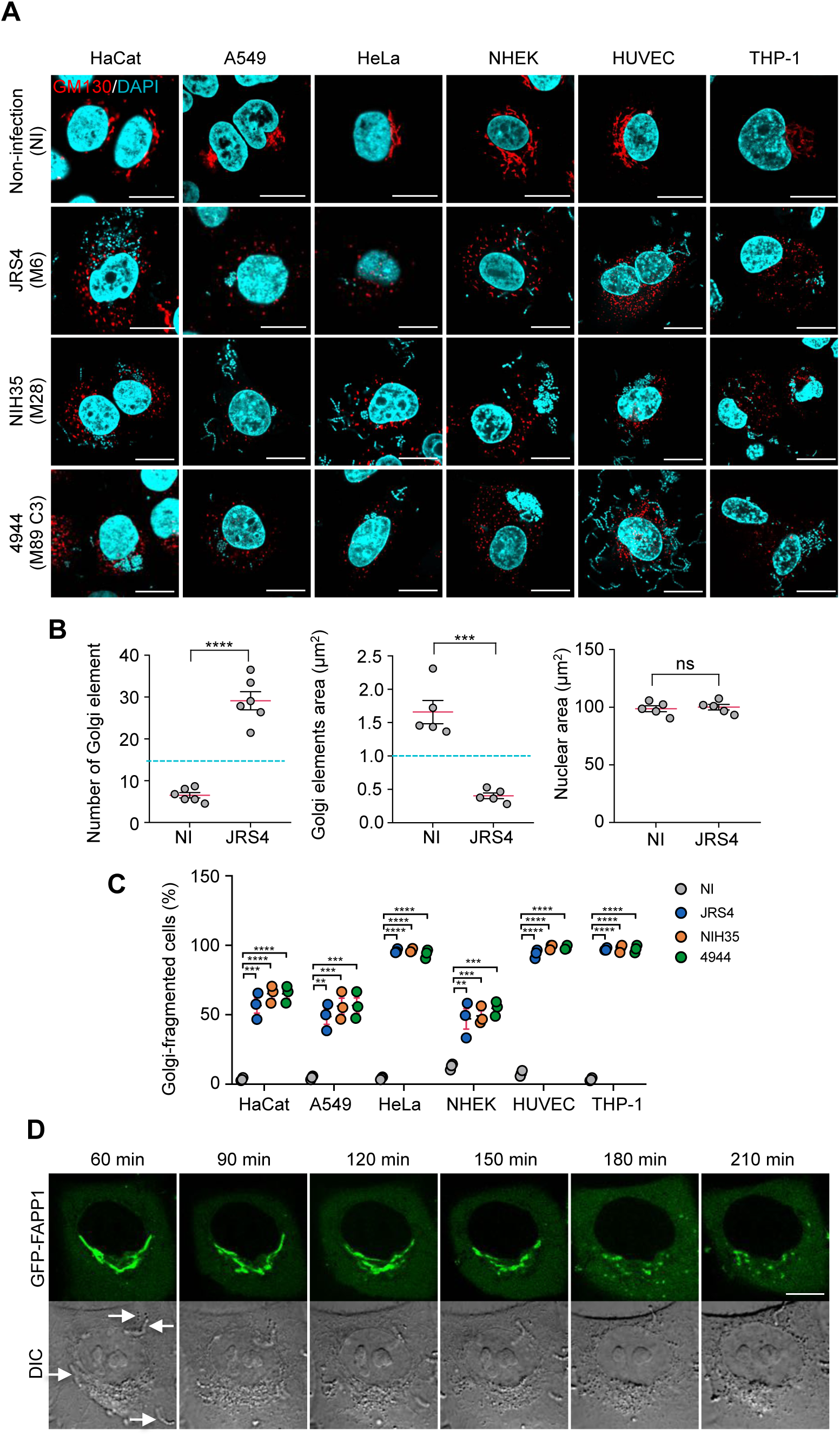
GAS induces Golgi fragmentation in infected host cells. (**A** and **B**) The Golgi structure during GAS infection. Cells were infected with indicated GAS strains for 4 h, fixed, and immunostained for the Golgi marker GM130 (red). Cellular and bacterial DNA DNA was stained with DAPI (cyan). Representative confocal images (A) and quantification of the Golgi and nucleus signals (**B**). (**C**) Quantification of the Golgi-fragmented cells that showed > 15 Golgi elements that is < 1 μm^2^. (**D**) EmGFP-FAPP1-expressing HeLa cells were infected with GAS JRS4. Confocal images were captured at indicated time after infection. Scale bars, 10 μm. Data in **B** and **C** represent individual values (dots) (n > 20 cells per condition) and the mean (magenta line) ± SEM of independent experiments. *P*-values calculated by two-tailed Student’s t-test. \*\**P* < 0.01, \*\*\**P* < 0.001, \*\*\*\**P* < 0.0001; ns, not significant.

To quantify the aforementioned phenotype, we measured the number and the area of Golgi fragments positive for the marker GM130; we found that the JRS4-infected cells harbored >20 Golgi elements, each with an area of 0.3 μm^2^–0.6 μm^2^ (Fig. 1B). We next defined cells containing >15 Golgi elements featuring an area of 1.0 μm^2^ as the Golgi-fragmented cells, and we found that the Golgi-fragmentation efficiencies were similar among the GAS strains (Fig. 1C); >90% of the infected HeLa cells, the HUVECs, and the THP-1 cells exhibited Golgi fragmentation at 4 h post-infection, whereas 40%–60% of the HaCat and the A549 cells and the NHEKs showed Golgi fragmentation (Fig. 1C).

To examine the time course of the changes in the Golgi apparatus structure during GAS infection, we expressed the EmGFP-tagged FAPP1 (a Golgi-resident protein) in cells and performed time-lapse imaging during the infection. In live-cell microscopy, the Golgi fragmentation process was detected at 2–3 h post-infection (Fig. 1D).

Golgi fragmentation has been reported in apoptotic cells [33]. Thus, to examine whether the infection-induced fragmentation here was caused by apoptotic signaling, we inhibited apoptotic signaling by overexpressing the antiapoptotic protein Bcl-2 [34]; ∼90% of the Bcl-2-expressing GAS-infected cells exhibited Golgi fragmentation (Fig. S2A). Moreover, this fragmentation was also not inhibited when infected cells were treated with the pan-caspase inhibitor Z-VAD-FMK (an inflammatory-caspase inhibitor) (Fig. S2B). These results suggest that apoptotic signaling may not be involved in the Golgi fragmentation that occurs during GAS infection. Collectively, our findings suggest that GAS infection trigger the fragmentation of the Golgi apparatus in various types of human cells.

### SLO and Nga are critical for GAS-induced Golgi fragmentation

Pathogenic bacteria inject virulence effector proteins into the host cells to modulate host cellular processes. GAS can deliver effector proteins across the host plasma membrane or the endosomal membrane to modulate host signaling by cytolysin-mediated translocation (CMT) that uses pore-forming cytolysin SLO. Therefore, to examine whether SLO functions in the infection-induced Golgi fragmentation described above, we infected the HeLa cells with a SLO-deficient mutant (Δ*slo*). Infection with JRS4 Δ*slo* did not induce Golgi fragmentation, whereas complementation with the gene *slo* completely rescued the phenotype (Fig. 2A and 2B). Moreover, JRS4 SLO^Y255A^, a mutant that lacks the pore-forming activity of SLO [20], failed to induce Golgi fragmentation (Fig. 2A and 2B), which indicates that GAS-induced Golgi fragmentation involves the pore-forming activity of SLO during infection.

**Fig. 2.**
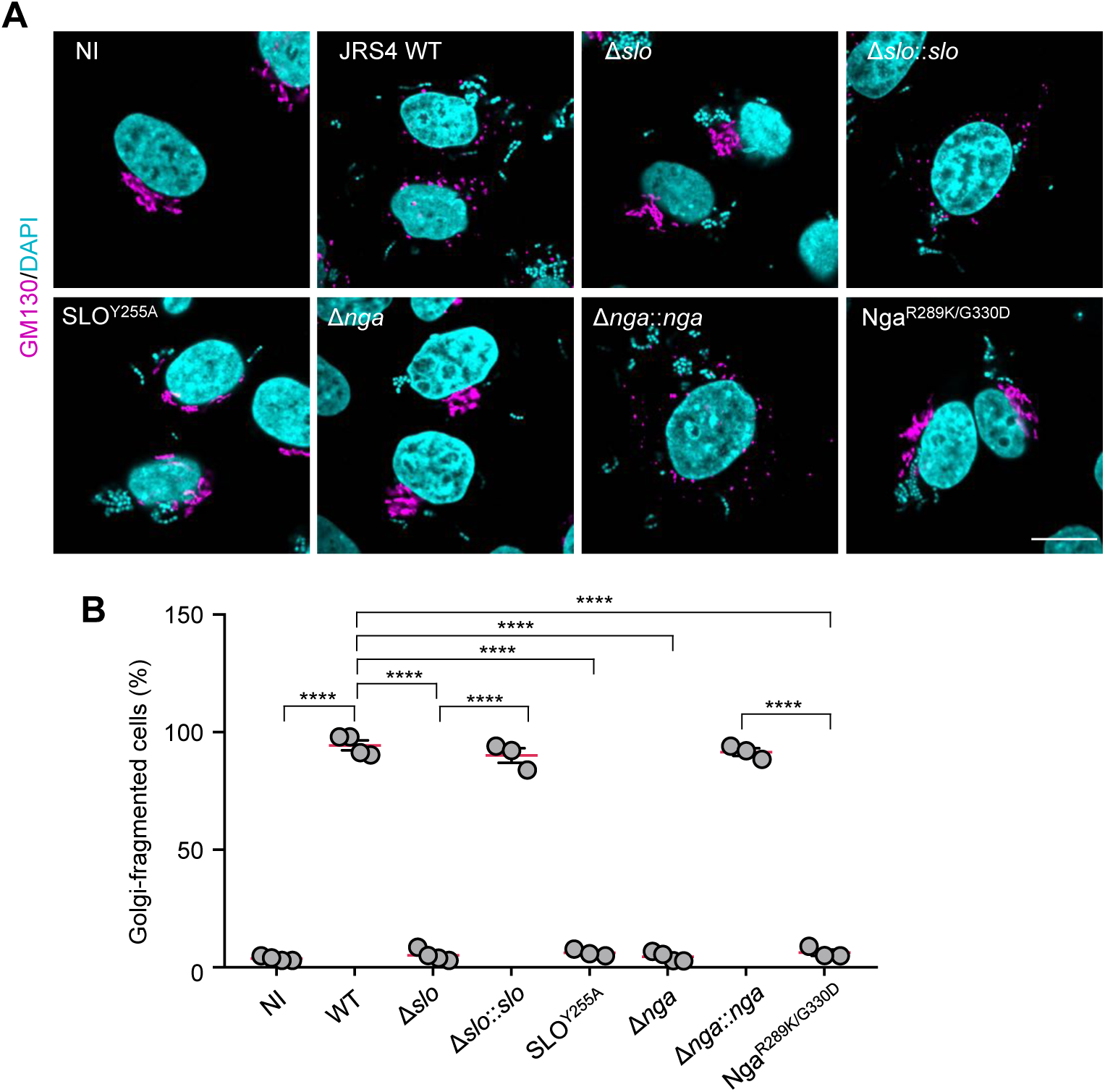
The Golgi fragmentation during GAS infection requires SLO and Nga. (**A** and **B**) HeLa cells were infected with indicated GAS strains for 4 h, fixed, and immunostained for GM130 (magenta). Representative confocal images (**A**), and quantification of the cells with fragmented Golgi during infection (**B**). Scale bars, 10 μm. Data in **B** represents individual values (dots) (n > 20 cells per condition) and the mean (magenta line) ± SEM of independent experiments. *P*-values calculated by two-tailed Student’s t-test. \*\*\*\**P* < 0.0001.

SLO is expressed from an operon that also encodes Nga, SLO is necessary for Nga translocation into the host cells [19]. To ascertain whether Nga was also required for GAS-induced Golgi fragmentation, or whether SLO pore-forming activity directly triggered the fragmentation, we examined the Golgi morphology in the cells infected with JRS4 Δ*nga* and Δ*nga*::*nga* (*nga*-complemented strain). The Golgi structures in the Δ*nga*-infected cells but not the Δ*nga*::*nga*-infected cells were found to be compact, and quantification of the Golgi signals indicated that *nga* was critical for GAS-induced Golgi fragmentation (Fig. 2A and 2B). We also confirmed that SLO and Nga were crucial for the Golgi fragmentation induced by the strain NIH35 (Fig. S3A and S3B).

To test whether the NADase activity of Nga was responsible for the Golgi fragmentation, we infected cells with the strain JRS4 Nga^R289K/G330D^; the mutations in Nga in the present study abolished the NADase activity of the effector [35]. Although JRS4 Nga^R289K/G330D^ can invade the host cytosol [23], we observed no alteration of the Golgi structure during JRS4 Nga^R289K/G330D^ infection (Fig. 2A and 2B). Taken together, these data indicate that SLO pore-forming activity and Nga NADase activity are required for GAS-induced Golgi fragmentation.

### GAS invasion is required for Nga-mediated Golgi fragmentation during infection

We next examined whether Golgi fragmentation was triggered by extracellular GAS. Because GAS JRS4 requires fibronectin-binding protein (FBP) for invading host cells [36], we constructed the strain JRS4 Δ*fbp* and infected the HeLa cells with this mutant; moreover, to monitor GAS invasion into the host cytosol, we expressed mCherry-Galectin-3, which serves as a marker of damaged vacuoles when invasive pathogens escape into the cytosol [37]. Our results confirmed that JRS4 Δ*fbp* were unable to invade the HeLa cells (Fig. 3A). Next, we tested whether JRS4 Δ*fbp* translocates Nga into the cytosol by analyzing NADase activity, assessed based on the NAD consumption, in the cytosol of the HeLa cells after infection. As hypothesized, after infection with the JRS4 wild-type and the Δ*fbp* strains, we measured comparable levels of NADase activity in the cytosol of the HeLa cells, which demonstrated that Nga was translocated across the host cell membrane even during infection with JRS4 Δ*fbp* (Fig. 3B). Unexpectedly, however, JRS4 Δ*fbp* failed to induce Golgi fragmentation (Fig. 3C). These results suggested that Golgi fragmentation require not only SLO secretion and Nga translocation into the cytosol, but also require GAS invasion into the host cells.

**Fig. 3.**
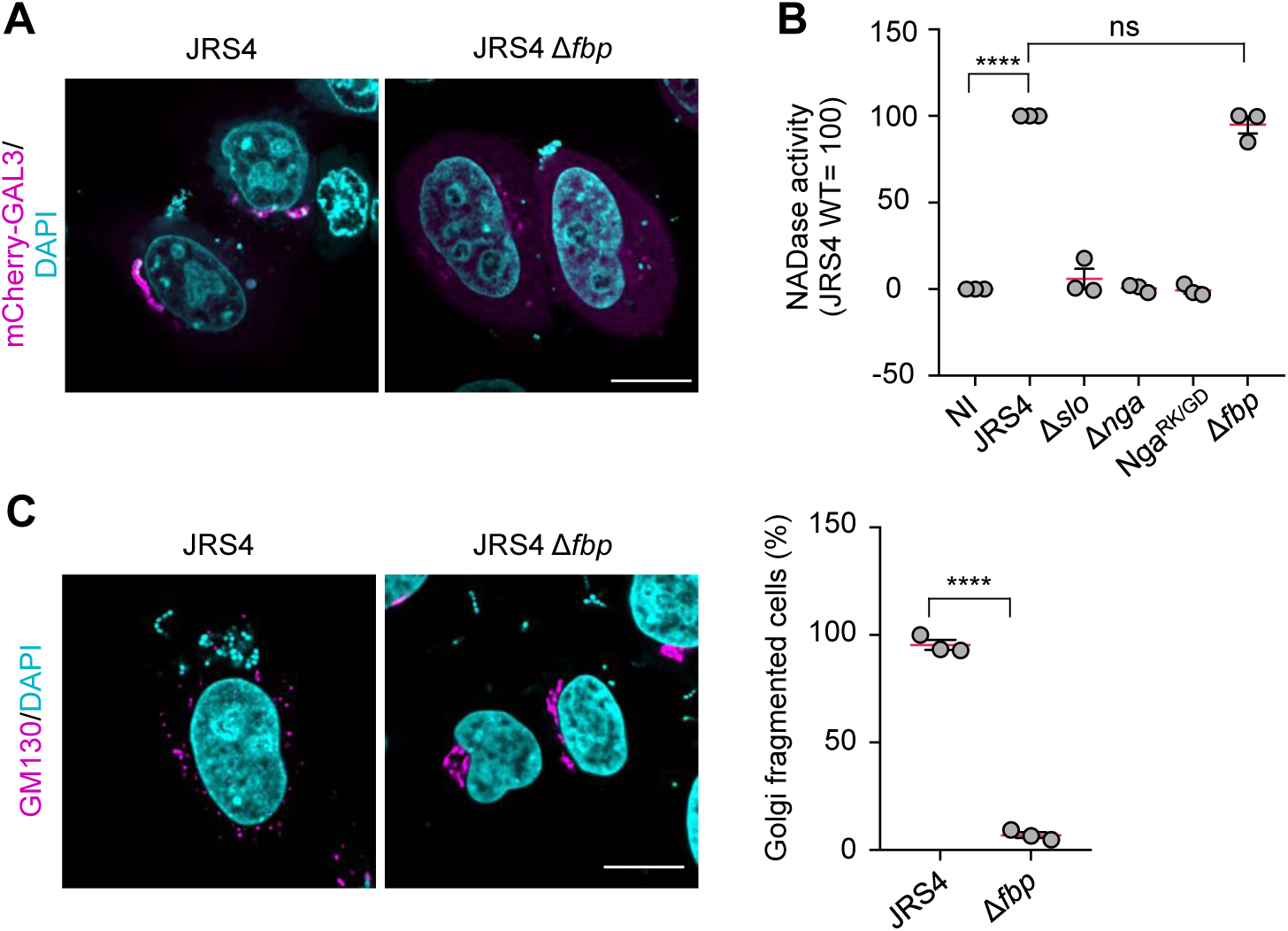
GAS invasion is necessary for Nga-mediated Golgi fragmentation. (**A**) HeLa cells transiently expressing mCherry-galectin-3 (GAL3) were infected with JRS4 wild-type or Δ*fbp* mutant for 4 h. mCherry-GAL3-positive Δ*fbp* was not observed. (**B**) NADase activity was assessed by measuring NAD consumption in the cytosolic fractions of infected cells. (**C**) HeLa cells were infected with indicated GAS strains for 4 h, fixed, and immunostained for GM130 (magenta). Scale bars, 10 μm. Data in **B** and **C** represent individual values (dots) (n > 20 cells per condition) and the mean (magenta line) ± SEM of independent experiments. *P*-values calculated by two-tailed Student’s t-test. \*\*\*\**P* < 0.0001; ns, not significant.

To exclude the possibility that FBP itself may be critical for the signaling that induces Nga-mediated Golgi fragmentation, we treated the cells with cytochalasin D (cytD) to inhibit GAS invasion; cytD treatment does not affect SLO-mediated translocation of Nga [21]. Notably, cytD treatment markedly suppressed Golgi fragmentation during JRS4 infection (Fig. S4). Together, these results showed that both GAS invasion and SLO-mediated injection of Nga into the host cells were critical for GAS-induced Golgi fragmentation.

### GAS impairs anterograde transport pathway

The Golgi apparatus functions in mediating protein and lipid modifications, transport, and sorting. To assess whether the post-Golgi secretion pathway was inhibited by GAS, we examined anterograde transport by using the RUSH (Retention Using Selective Hooks) system [38]. In our assay, E-cadherin was fused to a streptavidin-binding peptide (SBP) and EGFP and coexpressed with Streptavidin-KDEL, which localizes in the ER. Under normal conditions, interaction of the SBP-EGFP-E-cadherin with Streptavidin-KDEL in the ER prevented the transport of the fusion protein to the plasma membrane (Fig. S5). However, after the addition of biotin, which competes with the SBP tag for streptavidin binding, the SBP-EGFP-E-cadherin was released from the ER and transported to the plasma membrane through the Golgi complex in the non-infected cells, and the E-cadherin that was normally trafficked to the plasma membrane was detected by immunostaining for EGFP without membrane permeabilization (Fig. 4A). By contrast, in the JRS4-infected cells, the SBP-EGFP-E-cadherin exhibited punctate localization and the surface-EGFP signal was rarely detected (Fig. 4A). Quantification of the surface-EGFP signal revealed that anterograde trafficking of E-cadherin was abolished in the JRS4-infected cells (Fig. 4B). We also infected cells with Δ*slo*, Δ*nga*, Δ*nga*::*nga*, and Nga^R289K/G330D^ mutants, and we found that while Δ*slo*, Δ*nga*, and Nga^R289K/G330D^ did not affect anterograde trafficking, Δ*nga*::*nga* infection inhibited the trafficking as effectively as that by the JRS4 wild-type infection (Fig. 4A and 4B). Collectively, these results suggest that the Golgi fragmentation is caused by the invading GAS and the effector Nga results in the defect in the host-cell anterograde trafficking.

**Fig. 4.**
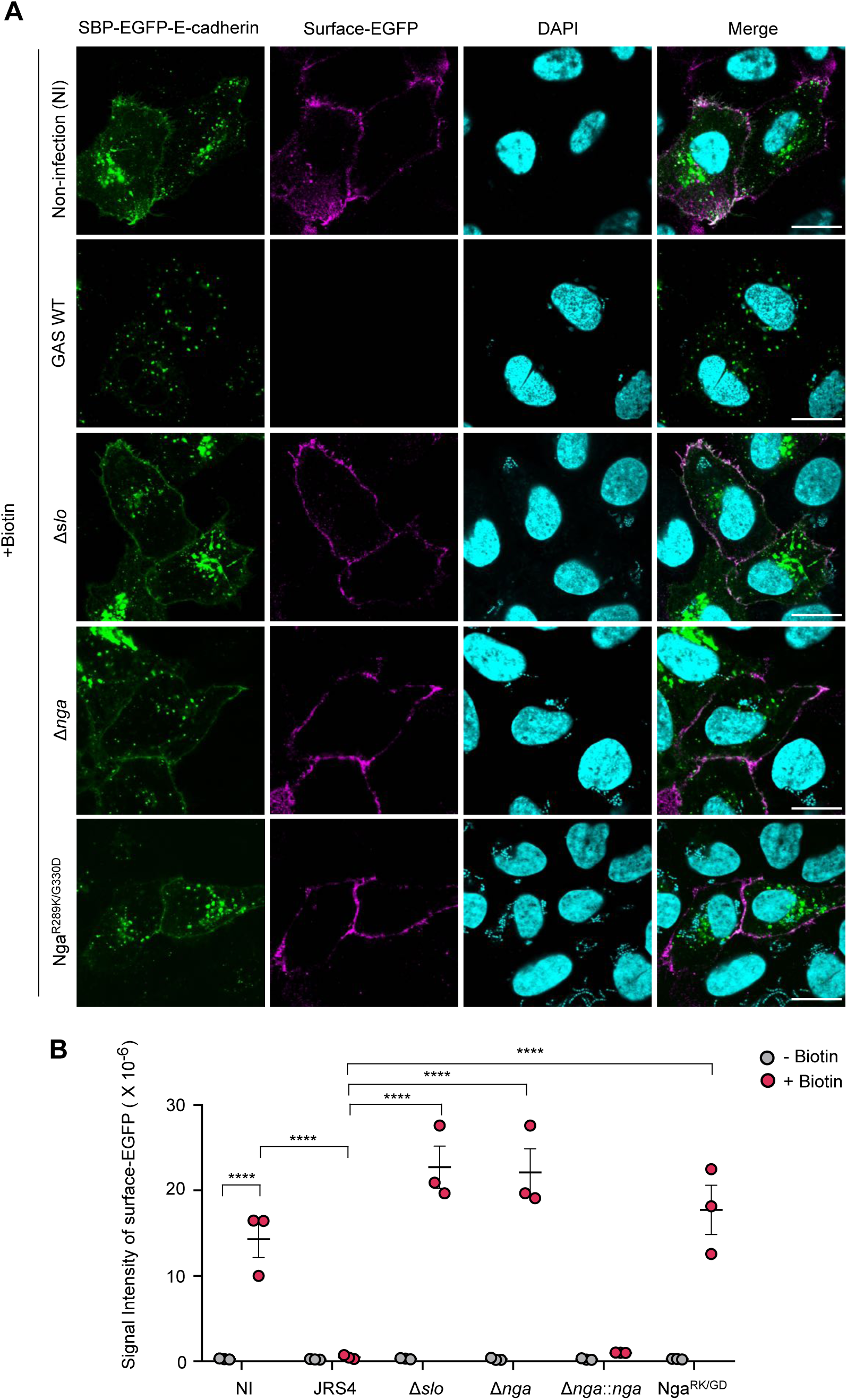
GAS inhibits anterograde transport through SLO and Nga. (**A**) Anterograde trafficking was inhibited in GAS infected cells. HeLa cells expressing Streptavidin-KDEL and SBP-EGFP-E-cadherin were infected with GAS strains. Cells were infected with GAS for 2 h, and then incubated for 1 h with biotin to observe the traffic of SBP-EGFP-E-cadherin to the plasma membrane. Cells were then fixed and the surface E-cadherin was detected with an anti-GFP (magenta) prior to cell permeabilization. Cellular and bacterial DNA DNA was stained with DAPI (cyan). Scale bars, 10 μm. (**B**) Quantification of surface E-cadherin using anti-GFP immunostaining. Average intensity of regions of interest corresponding to transfected cells was measured. Data in **B** represents individual values (dots) (n > 20 cells per condition) and the mean (magenta line) ± SEM of independent experiments. *P*-values calculated by two-tailed Student’s t-test. \*\*\*\**P* < 0.0001; ns, not significant.

### Invading GAS and effector Nga disrupt epithelial integrity

E-cadherin is critical for the cell-cell adhesion that holds epithelial cells tightly together, and thus E-cadherin is a crucial molecule for the maintenance of the epithelial barrier. GAS can translocate across epithelial barriers by degrading junctional proteins, including E-cadherin [39]. Therefore, we examined whether the inhibition of anterograde trafficking by Nga affects E-cadherin localization and the ability of GAS to translocate across epithelial monolayers. Because the GAS protease SpeB degrades E-cadherin, we used a JRS4 strain that was defective in SpeB expression. Immunostaining of the HaCat cells revealed that while E-cadherin was confined to the cell membrane in the non-infected cells and in cells infected with the strain Nga^R289K/G330D^, E-cadherin was present in substantial amounts in the cytoplasm in the JRS4-infected cells (Fig. 5A); this result suggests that the trafficking of endogenous E-cadherin to the cell membrane may be impaired by the Nga derived from the invading GAS. Furthermore, the HaCat cells treated with brefeldin A (BFA), which inhibits ARF and induces Golgi fragmentation [40], also exhibited E-cadherin redistribution similar to that induced by a JRS4 infection (Fig. 5A). We also examined the total E-cadherin level in cells and found that JRS4 infection did not affect the cellular E-cadherin amounts (Fig. 5B). These results suggest that Nga may not degrade E-cadherin but may alter the subcellular localization of E-cadherin during GAS infection.

**Fig. 5.**
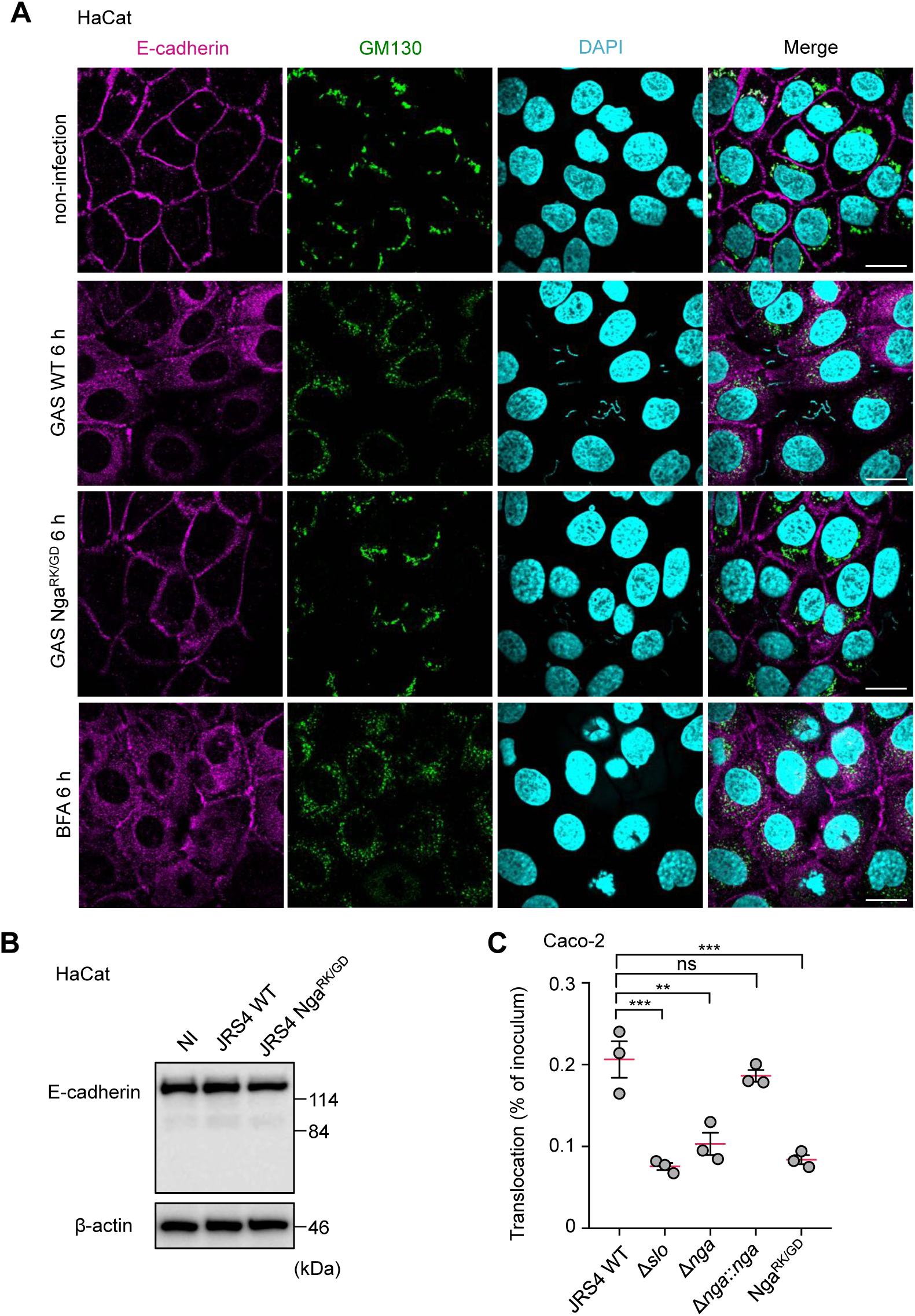
GAS affects E-cadherin trafficking and translocation of GAS through Nga. (**A**) HaCat cells were infected with GAS strains for treated with BFA for 6 h, fixed and immunostained with anti-E-cadherin (green) and GM130 (magenta). Cellular and bacterial DNA DNA was stained with DAPI (cyan). Scale bars, 10 μm. (**B**) Western blotting of indicated proteins in GAS-infected HaCat cells (6 h). (**C**) Caco-2 cells were grown on Millicell filters and then infected with GAS strains at an MOI of 10 for 6 h. Bacterial translocation was expressed as a percentage of GAS recovered from medium beneath the monolayer at 6 h after infection. Data in **C** represents individual values (dots) and the mean (magenta line) ± SEM of independent experiments. *P*-values calculated by two-tailed Student’s t-test. \*\**P* < 0.01, \*\*\**P* < 0.001; ns, not significant.

Redistribution of E-cadherin in the GAS-infected cells may increase bacterial translocation through the paracellular pathway. Because the HaCat, the A549, and the HeLa cells exhibit unstable junctional integrity, as indicated by their measured transepithelial electrical resistance (TER) [41], these epithelial cells are not suitable for assessing GAS translocation. Thus, for assaying GAS translocation, we selected the polarized Caco-2 cells, which are widely used as an *in vitro* model of the epithelial barrier and have been previously used in experiments on GAS translocation [39,42]. We confirmed that GAS infection induced Golgi fragmentation even in Caco-2 monolayers (Fig. S6). The apical surface of the Caco-2 monolayers was infected with either the JRS4 wild-type or the JRS4 Δ*slo*, Δ*nga*, Δ*nga*::*nga*, or Nga^R289K/G330D^ mutant for 1 h, and the bacteriostatic agent trimethoprim was added to inhibit the additional growth of extracellular bacteria and translocated bacteria. At 6 h post-infection, translocated bacteria were examined using the colony formation assay. Relative to the JRS4 wild-type, the mutants Δ*slo*, Δ*nga*, and Nga^R289K/G330D^ exhibited markedly diminished translocation efficiency, whereas Δ*nga*::*nga* showed comparable translocation efficiency (Fig. 5C). These results suggest that SLO and Nga facilitate GAS translocation across epithelial monolayers, perhaps by disrupting intracellular trafficking.

### Invading GAS inhibits IL-8 secretion by using Nga

GAS infection-induced Golgi fragmentation was also observed in the differentiated THP-1cells (Fig. 1A), and this fragmentation occurred through an Nga-dependent mechanism (Fig. 6A). Because macrophages produce the chemokine IL-8 in response to bacterial infection, we next determined whether invading GAS inhibited IL-8 secretion by using Nga. The differentiated THP-1 cells secreted IL-8 in response to infection by Δ*slo*, Δ*nga*, and Nga^R289K/G330D^ but not JRS4 wild-type or Δ*nga*::*nga* (Fig. 6B), which suggests that SLO and Nga may inhibit IL-8 secretion by macrophages. Moreover, the invasive GAS strain NIH35 blocked IL-8 secretion through an SLO- and an Nga-dependent mechanism (Fig. 6B). To exclude the possibility that the lack of IL-8 production may be due to the suppression of IL-8 expression by SLO and Nga during GAS infection, we examined IL-8 secreted from the LPS-primed macrophages. The LPS-induced secretion of IL-8 was also inhibited by SLO and Nga (Fig. 6C), which indicates that the IL-8 secretion process was blocked by Nga during GAS infection.

**Fig. 6.**
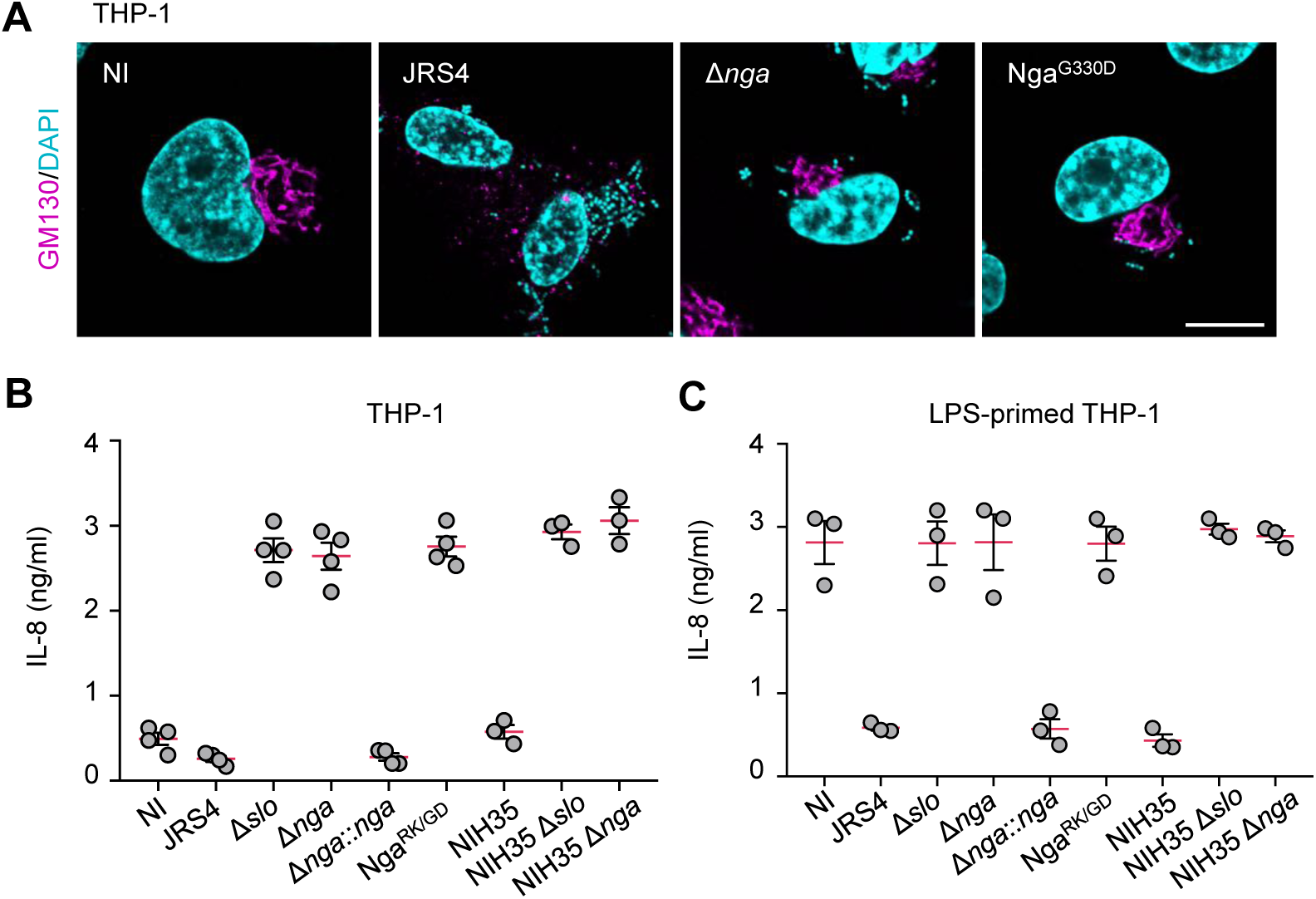
GAS inhibits IL-8 secretion process through SLO and Nga. (**A**) Differentiated THP-1 cells were infected with GAS JRS4 strains for 4 h, fixed, and immunostained with GM130 (magenta). Cellular and bacterial DNA was stained with DAPI (cyan). Scale bars, 10 μm. (**B** and **C**) Non-primed (**B**) or LPS-primed (**C**) differentiated THP-1 cells were infected with GAS strains for 4 h. Supernatants were analyzed for the secretion of IL-8 by ELISA.

## Discussion

Within bacterium-infected cells, highly complex interactions occur between the host immune-system components and the bacterial pathogen, and unique molecular dynamics are frequently observed [25]. We discovered in the present study that GAS invasion induced the fragmentation of the Golgi complex and inhibited the anterograde transport in an SLO- and an Nga-dependent manner. Notably, although GAS was found to translocate Nga into the host cytosol through an SLO-dependent mechanism without invading the host cell, a noninvasive GAS mutant (JRS4 Δ*fbp*) did not trigger Golgi fragmentation. These results uncovered a previously unknown function of Nga that this effector protein performs in conjunction with other effectors and/or the GAS invasion process.

To our knowledge, this is the first report that GAS invasion disrupts the Golgi complex and the post-Golgi secretory pathway. The Golgi complex functions in sorting and trafficking in the central vacuolar system, and the Golgi apparatus and Golgi-associated trafficking have been widely reported to be affected by bacterial infection [25]. For example, during *Shigella* infection, the *Shigella* effector protein IpaB induces cholesterol relocation and disrupts the Golgi complex and anterograde and retrograde transport [30]; these modifications lead to the disruption of the host epithelial barrier and are associated with *Shigella* pathogenesis. We showed in the present study that GAS Nga activity also inhibited E-cadherin trafficking to the plasma membrane. E-cadherin promoted the cell-to-cell adhesion and integrity of the epithelial barrier, and, accordingly, GAS translocation across epithelial monolayers was suppressed by the knockout of *slo* or *nga*. Sumitomo et al have reported that streptolysin S and a cysteine protease contribute to bacterial translocation by perhaps directly destabilizing intercellular junction proteins such as E-cadherin [39,42,44]. Our data suggest that invading GAS may support the translocation of extracellular GAS and may facilitate invasion into deeper tissues.

Intriguingly, VirA from *S. flexneri* and EspG from enteropathogenic *E. coli* directly inactivate Rab1 and disrupt ER-to-Golgi trafficking in cells, and this disruption of the host secretory pathway results in the inhibition of IL-8 secretion from the infected cells [31]; this suggests that the impairment of the post-Golgi secretory pathway may be linked to the attenuation of the inflammatory response. Lethal necrotizing fasciitis caused by GAS is characterized by the presence of few neutrophils at the infection site, and GAS expresses a secretory protein that degrades IL-8, which is crucial for neutrophil transmigration and activation [45-49]. Thus, the absence of anterograde transport in GAS-invaded cells likely contributes to the GAS pathogenesis. Recently, newly emergent clade-associated strains of serotype M89 (M89 clade-3 strains) continue to be recognized as a cause of invasive diseases worldwide, and these strains were found to be genetically acapsular and thus incapable of producing the hyaluronic acid capsule [13,50]. Because the hyaluronic acid capsule is a critical virulence factor required for evading phagocytosis or endocytosis by host cells [51,52],dissemination of these strains may be associated with the ability to invade host cells; however, the precise mechanism by which the acapsular characteristics influence the pathogenesis of these strains remains unknown. Although the expression of SLO and Nga is enhanced in M89 clade-3 strains, clade-associated and non-clade-associated M89 strains exhibit comparable intracellular survival [13,50]. Therefore, a previously unrecognized function of the Nga derived from intracellular GAS that suppresses host immune responses may be associated with the pathogenicity of the M89 clade-3 strains.

GAS invasion and the effector Nga are visibly linked to the morphological and the functional destruction of the Golgi complex, but the molecular mechanism underlying this process has remained unclear. Unexpectedly, we found that the GAS JRS4 Δ*fbp*, which can inject Nga into the host cytosol, did not induce Golgi fragmentation; this suggested that Nga alone may be insufficient for inducing the fragmentation. Although the proteins and/or events that function in Nga-dependent Golgi fragmentation during GAS invasion remain to be identified, the fragmentation was observed in all GAS strains tested, which indicates that certain common characteristics shared among the strains are involved in producing this phenotype. Our time-lapse imaging analysis revealed that Golgi fragmentation occurred starting from 2 h–3 h post-infection, which coincides with the time of GAS invasion into the cytoplasm. Therefore, we hypothesize that the unidentified molecules secreted from GAS may be involved in the Golgi fragmentation.

In summary, our findings indicate that GAS infection disrupts the Golgi-related network in host cells through the effector Nga and intracellular GAS, which then enables the translocation of GAS across epithelial barriers and the inhibition of IL-8 secretion by macrophages *in vitro*. Further investigation aimed at identifying other GAS molecules responsible for the Golgi fragmentation will enhance our understanding of the pathogenicity of GAS.

## Methods

### Bacterial strains and infection

GAS strains JRS4, NIH35, and 4944 were grown in Todd-Hewitt broth supplemented with 0.2% yeast extract (THY) at 37°C. The isogenic mutant strains JRS4 Δ*slo*, JRS4 Δ*nga*, JRS4 Δ*nga*::*nga*, and JRS4 Nga^R289K/G330D^ have been described previously [53]. JRS4 Δ*fbp*, NIH35 Δ*slo*, and NIH35 Δ*nga* were constructed using a two-step allele exchange by a method described previously. Overnight cultures were reinoculated in fresh THY and grown to the exponential phase (OD600: 0.7–0.8), collected by centrifugation, and diluted with cell culture media before use. Cell cultures in media without antibiotics were infected for 1 h with GAS at a multiplicity of infection (MOI) of 100. Infected cells were washed with phosphate-buffered saline (PBS) and treated with 100 μg/mL gentamicin for an appropriate period to kill the bacteria that were not internalized.

### Cell culture

HeLa, A549, and THP-1 cell lines were purchased from ATCC; HaCat cell line was a gift from Dr. Kabashima; HUVECs and NHEKs were purchased from PromoCell; and Caco-2 cells were purchased from the Riken Cell Bank. HeLa and A549 cells were maintained in the Dulbecco’s modified Eagle’s medium (Nacalai Tesque) supplemented with 10% fetal bovine serum (FBS; Gibco) and 50 μg/mL gentamicin (Nacalai Tesque), and the THP-1 cells were cultured in RPMI 1640 medium (Nacalai Tesque) supplemented with 10% FBS and 50 μg/mL gentamicin. THP-1 cells were differentiated into macrophages by stimulating them with 50 ng/mL phorbol 12-myristate for 72 h. HUVECs were maintained in Endothelial Cell Growth Medium 2 Kit (PromoCell) supplemented with 10% FBS and 50 μg/mL gentamicin; NHEKs were cultured in Keratinocyte Growth Medium 2 Kit (PromoCell); and Caco-2 cells were maintained in minimum essential medium (Wako) supplemented with 10% FBS and 50 μg/mL gentamicin. Cells were incubated in a 5% CO2 incubator at 37°C.

### Fluorescence microscopy

Immunofluorescence analysis was performed using these antibodies: anti-TOMM20 (ab78547; Abcam, 1:100), anti-calnexin (610524; BD Transduction Laboratories, 1:100), anti-GM130 (610822; BD Transduction Laboratories, 1:100), anti-TGN46 (13573-1-AP; Proteintech, 1:100), anti-GFP (GF200; 04363-24; Nacalai Tesque, 1:100), anti-E-cadherin (24E10; 3195; Cell Signaling Technology, 1:100), and anti-TOM20 (F-10; sc-17764; Santa Cruz Biotechnology, 1:100). The secondary antibodies used were anti-mouse or anti-rabbit IgG conjugated to Alexa Fluor 488 or 594 (#A32723, #A32742, #A32731, #A32740; Invitrogen). Cells were washed with PBS, fixed with 4% paraformaldehyde (PFA) in PBS (15 min), permeabilized with 0.1% Triton in PBS (10 min), washed with PBS, and blocked (room temperature, 1 h) with a skim milk solution (5% skim milk, 2.5% goat serum, 2.5% donkey serum, and 0.1% gelatin in PBS) or a BSA solution (2% BSA and 0.02% sodium azide in PBS). Next, the cells were probed (room temperature, 1 h) with the primary antibodies diluted in a blocking solution, washed with PBS, and labeled with the appropriate secondary antibody. To visualize bacterial and cellular DNA, cells were stained with4′,6-diamidino-2-phenylindole (DAPI; Dojindo). Confocal fluorescence micrographs were acquired using an FV1000 laser-scanning microscope (Olympus).

### Plasmids, transfection, and reagents

Human FAPP1 cDNA was PCR-amplified from the HeLa cell total mRNA and cloned into pcDNA-6.2/N-EmGFP-DEST (for N-terminal tagging) by using the Gateway (Invitrogen) cloning technology as described previously [54]. Str-KDEL_TNF-SBP-EGFP was purchased from Addgene (Plasmid #65278). Polyethylenimine (Polysciences) and Lipofectamine 3000 (Invitrogen) were used for transfection. Z-VAD-FMK was purchased from Promega.

### Measurement of NADase activity

HeLa cells seeded at 1.5 × 10^5^ cells/well in 24-well plates were infected with the GAS strains for 3 h without killing of extracellular bacteria by gentamicin. Infected HeLa cells were scrapped with chilled PBS, pooled with debris, and lysed in sterile water. The whole-cell lysates were cleared from the membrane fraction by centrifugation at 20,300 × *g* for 30 min at4°C to obtain the cytosolic fraction, which was diluted 2-fold with 2× PBS. NAD^+^ (Nacalai Tesque) was added to the cytosolic fraction at 1 mM and the mixtures were incubated at 37°C for 3 h. To develop reactions, NaOH (5 N) was added to the reaction mixtures, which were then incubated in the dark at room temperature for 30 min. Samples were analyzed by using a Wallac ARVO SX Multilabel Counter (Perkin Elmer) at 340-nm excitation/460-nm emission to examine the fluorescence intensity of the remaining NAD^+^. NAD^+^ hydrolysis levels in lysates from the wild-type GAS-infected cells and the non-infected cells were set to correspond to 100% and 0% NADase activity, respectively.

### RUSH assay

To assess anterograde transport, the HeLa cells were transfected with the RUSH plasmid (Streptavidin-KDEL_E-cadherin-SBP-EGFP), and at 24 h post-transfection, cells were infected with GAS strains as described above. After infection for 2 h, 40 μM biotin (Nacalai Tesque) was added to the cells, and after incubation for 1 h, the cells were fixed with 4% PFA (15 min) and immunostained with anti-GFP antibody without permeabilization. Images were acquired using confocal microscopy and analyzed using an ImageJ software. Regions corresponding to the transfected cells were drawn and the average intensity of surface anti-GFP staining in these regions was determined; >20 cells were analyzed under each condition and three independent experiments were performed.

### Translocation assay

Caco-2 cells were seeded at 2 × 10^5^ cells/well onto polycarbonate Millicell culture-plate inserts (12-mm diameter, 3-μm pore size; Millipore) and cultured for 5 days. To determine the Caco-2-cell monolayer integrity, the TER of the monolayers on the filter was measured using a Millicell-ERS device (Millipore), and monolayers with measured TER values of 450–500 Ω/cm^2^ were used in experiments. After the polarized monolayers were infected for 1 h with GAS (MOI = 10), the non-adherent bacteria were removed by washing the upper chamber with PBS, and the medium was switched to medium containing trimethoprim to inhibit the additional growth of GAS. The ability of GAS to translocate across monolayers was assessed through quantitative culturing of the medium: Each medium sample obtained from the lower chamber at 6 h post-infection was serially diluted and plated on THY agar plates to determine colony forming unit (CFU) values.

### Chemokine and cytokine secretion

THP-1 cells were seeded at 2 × 10^5^ cells/well in 24-well plates and differentiated for 72 h. After infection with GAS for 4 h, the supernatant was collected and centrifuged, and the IL-8 released into the supernatant was quantified by using a human IL-8 ELISA kit (Proteintech) according to manufacturer instructions.

### Statistical analysis

Values, including plotted values, represent means ± standard error of the mean (SEM). Data were tested using the two-tailed Student’s *t*-test, and *P* < 0.05 was considered significant (\**P*< 0.05, \*\**P* < 0.01, \*\*\**P* < 0.001, \*\*\*\**P* < 0.0001; ns, not significant). GraphPad Prism 8 was used for statistical analyses.

## Acknowledgments

This work was financial support from Grants-in-Aid for Scientific Research (16H05188, 15K15130, 26462776, 17K19552), the Takeda Science Foundation, the Research Program on Emerging and Re-emerging Infectious Diseases (18fk0108044h0202, 18fk0108073h0001) and J-PRIDE (18fm0208030h0002) from Japan Agency for Medical Research and Development, AMED.

## Figure legends

**Supplementary Fig. 1.**
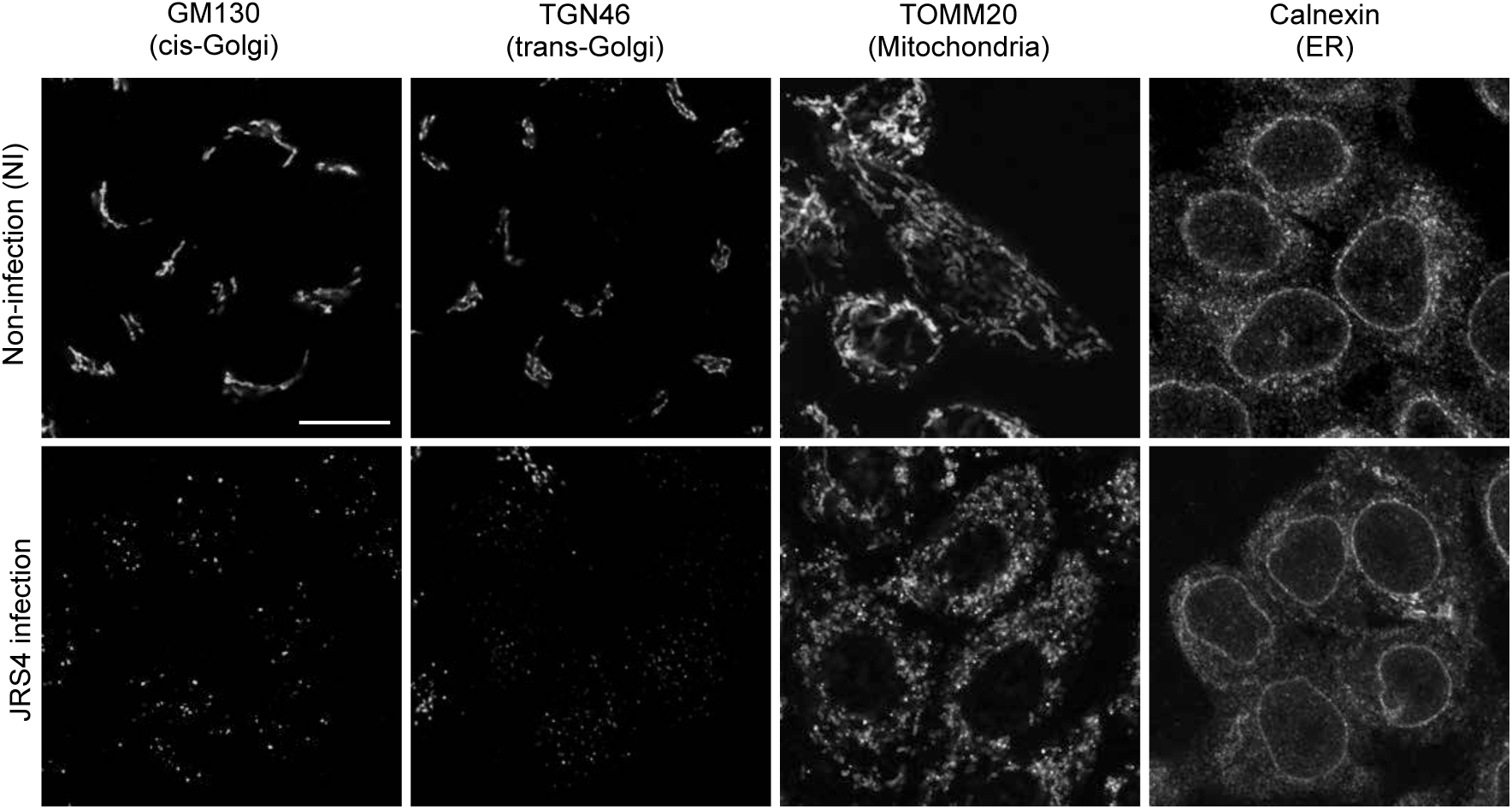
GAS infection induces the fragmentation of the Golgi and mitochondria. HeLa cells were infected with GAS JRS4 for 4 h, fixed, and immunostained with indicated antibodies.

**Supplementary Fig. 2.**
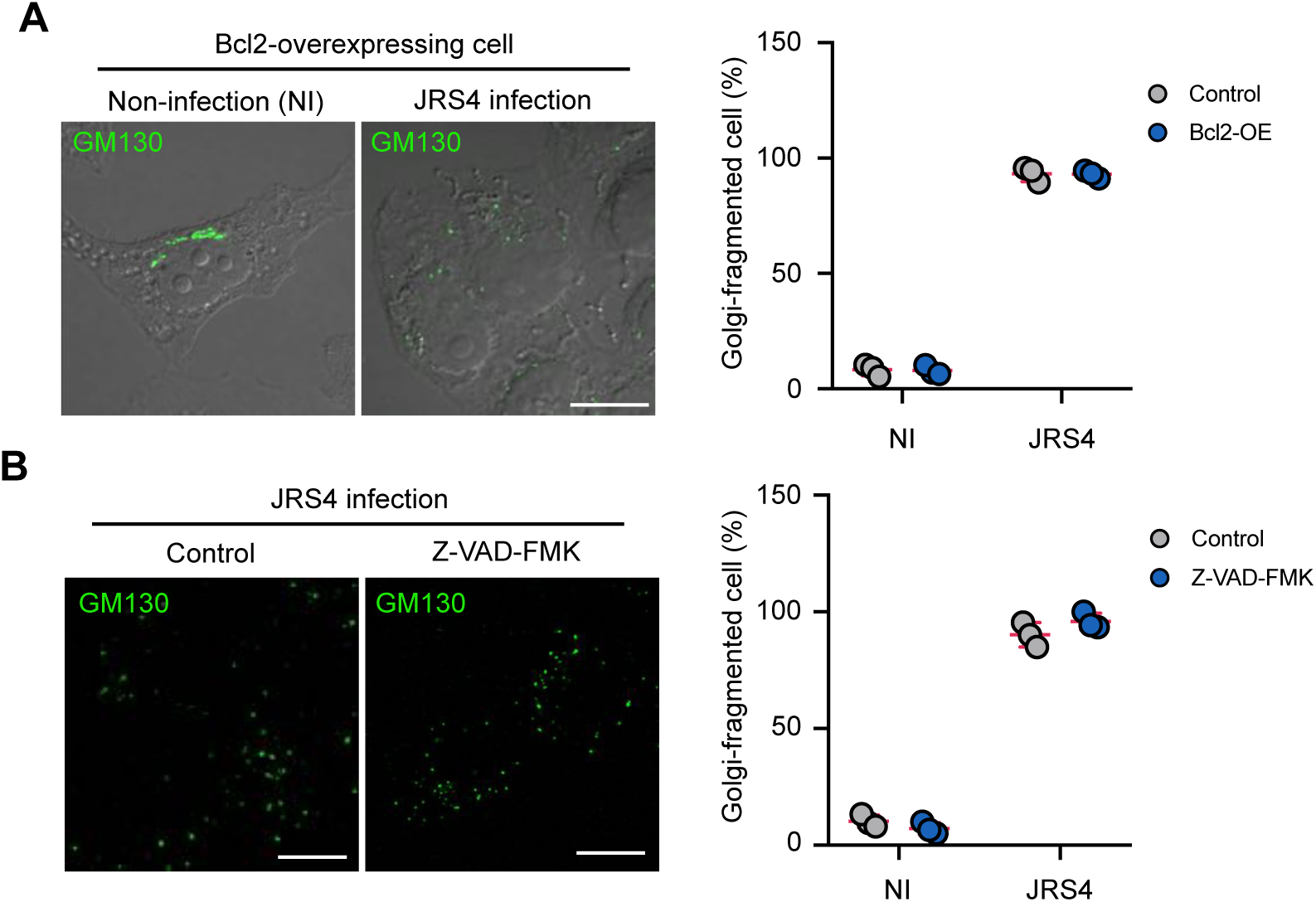
Inhibition of apoptotic signal does not suppress the Golgi fragmentation during GAS infection. HeLa Bcl-2-expressing cells were infected with GAS JRS4 for 4 h, and immunostained for GM130. The percentages of cells showing Golgi fragmentation were quantified. Data in **C** and **E** represent individual values (dots) (n > 20 cells per condition) and the mean (magenta line) ± SEM of independent experiments.

**Supplementary Fig. 3.**
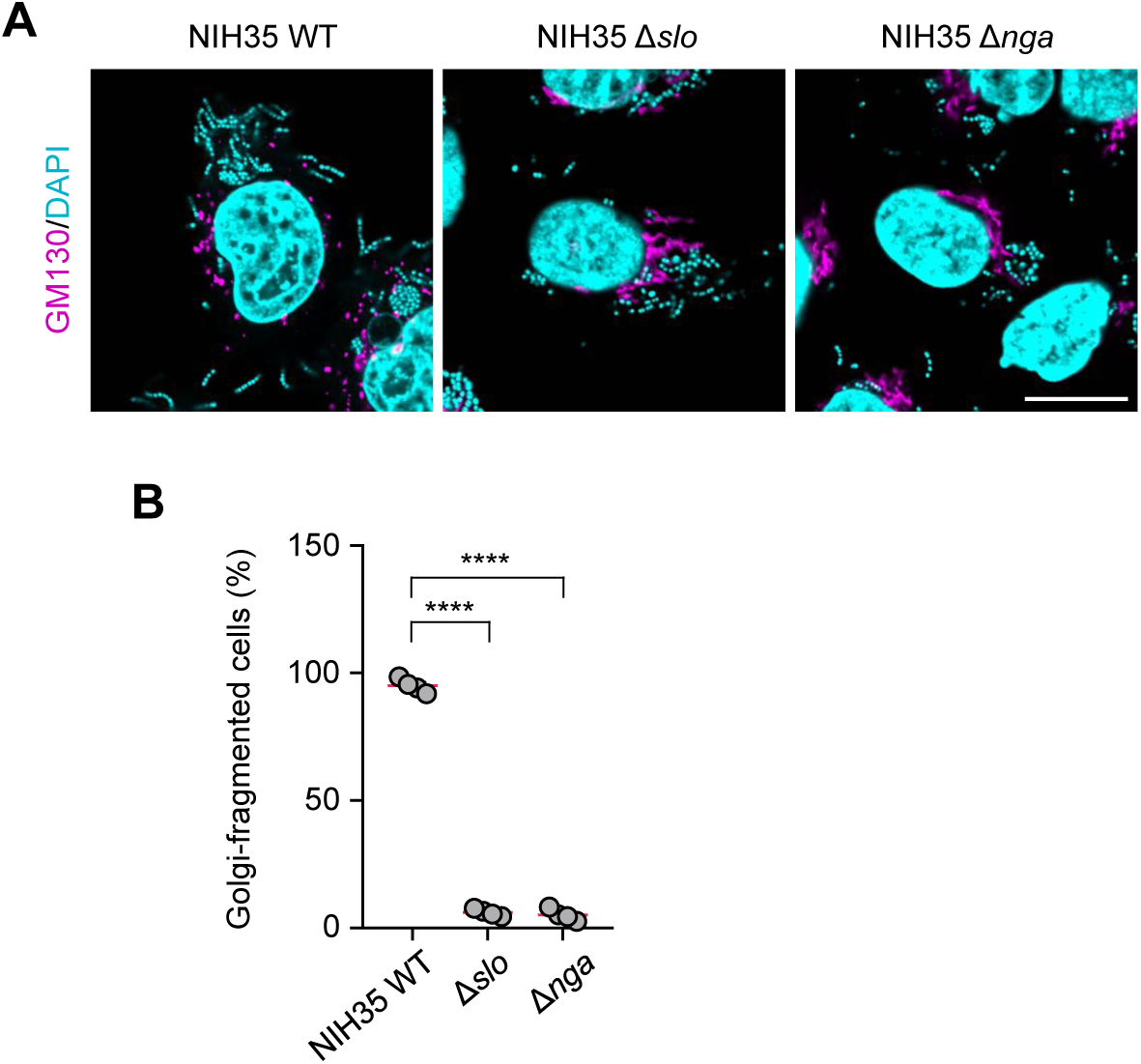
GAS NIH35 strain induces the Golgi fragmentation in a SLO- and Nga-dependent manner. HeLa cells were infected with NIH35 strains for 4 h, and immunostained for GM130. (**B**) The percentages of cells showing Golgi fragmentation were quantified. Data in **B** represents individual values (dots) (n > 20 cells per condition) and the mean (magenta line) ± SEM of independent experiments.

**Supplementary Fig. 4.**
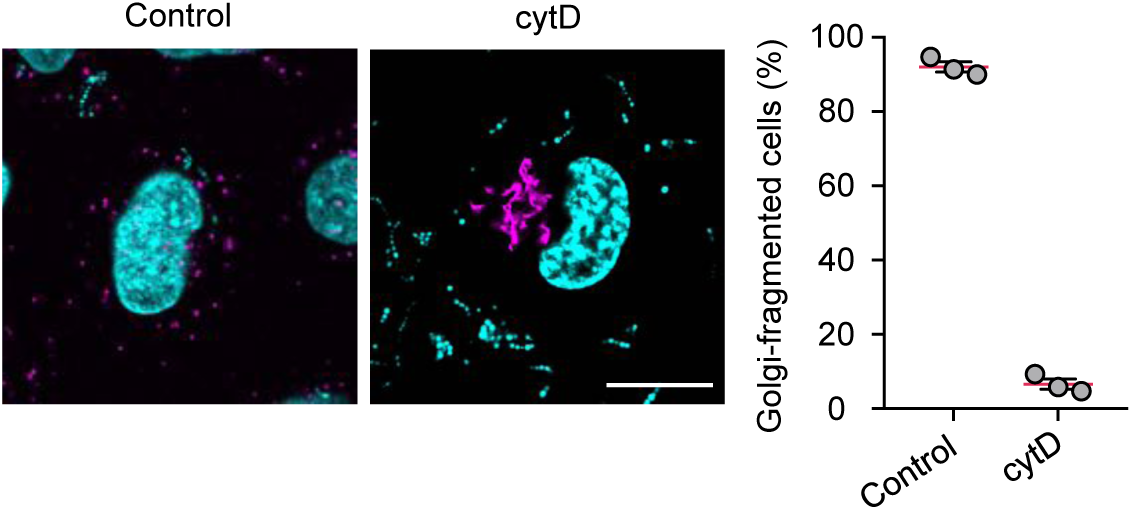
GAS NIH35 strain induces the Golgi fragmentation in a SLO- and Nga-dependent manner. HeLa cells treated with cytD were infected with GAS JRS4 for 3 h. Cells were immunostained for GM130. The percentages of cells showing Golgi fragmentation were quantified. Data represent individual values (dots) (n > 20 cells per condition) and the mean (magenta line) ± SEM of independent experiments.

**Supplementary Fig. 5.**
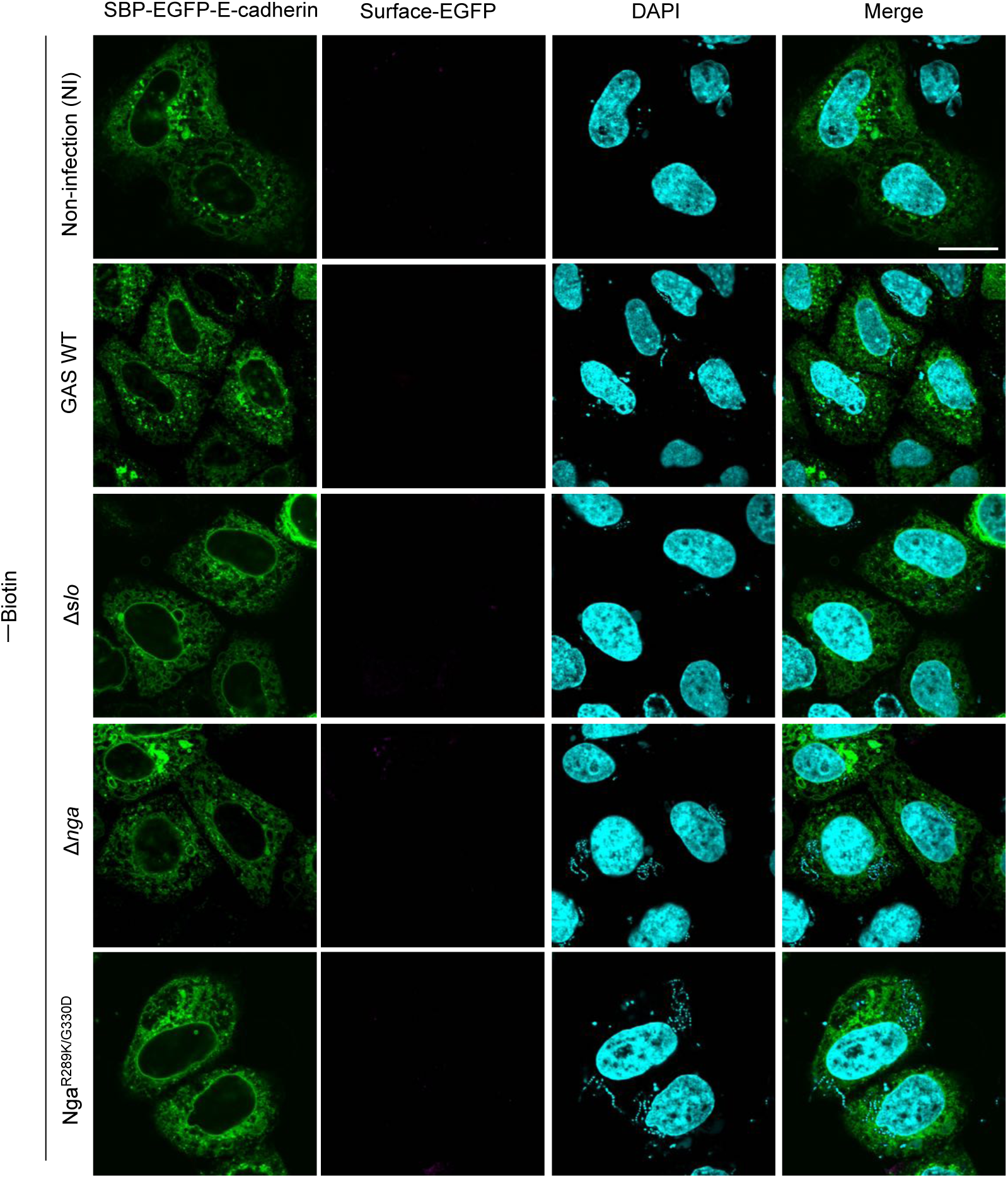
Confocal images of control condition in RUSH assay. HeLa cells expressing Streptavidin-KDEL and SBP-EGFP-E-cadherin were infected with GAS strains. Cells were then fixed and the surface E-cadherin was detected with an anti-GFP (magenta) prior to cell permeabilization. Cellular and bacterial DNA was stained with DAPI (cyan). Scale bars, 10 μm.

**Supplementary Fig. 6.**
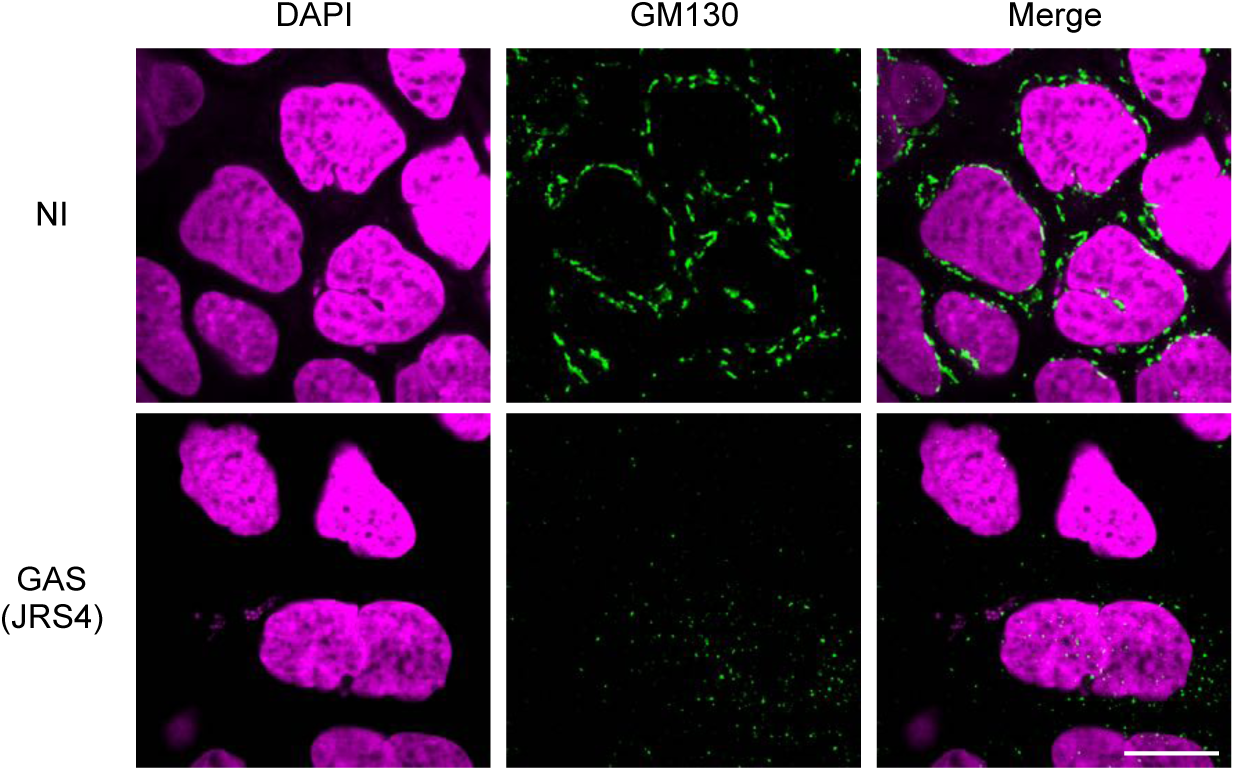
GAS infection induced the Golgi fragmentation in Caco-2 cells. Caco-2 cells were non-infected or infected with JRS4 GAS for 4 h, and immunostained for GM130. Scale bars, 10 μm.

